# Photo-based individual identification is more reliable than visible implant elastomer tags or toe-tipping in young agile frogs

**DOI:** 10.1101/2025.06.19.660530

**Authors:** Edina Nemesházi, Nikolett Ujhegyi, Zsanett Mikó, Andrea Kásler, Vera Lente, Veronika Bókony

## Abstract

In amphibian capture-recapture studies, commonly used individual-identification methods include toe clipping as well as less invasive alternatives of varying cost. Yet, choosing the best method for a study is challenging, because both the reliability for identification and the severity of adverse effects of a given mark type can greatly vary between species as well as life stages. We compared the reliability of three identification methods in young agile frogs (*Rana dalmatina*): clipping a single phalanx, injecting visible-implant-elastomer (VIE) tags (one of six colours) under the skin, and photo-based identification using natural colouration. Individuals were regularly photographed from the start of metamorphosis onwards, and were marked by the other two methods soon after all of them finished metamorphosis. Subsequently, we checked mark retention by each method multiple times for more than a year. Photo-based identification was far more reliable than the other two methods: 100% identification success post-metamorphosis in all checking events within small housing groups, and 98% with computer-assisted identification across all housing groups. Post-metamorphic body colouration remained largely stable, and the major patterns were already present at metamorphosis. Based on VIE colour alone, we could correctly identify the animals in 77% of the checks. Green tags were the easiest to recognize. VIE often broke into smaller parts, and those were sometimes found only outside the originally tagged body part (15%) or not found at all (10%). VIE was better retained in the legs than in the arms. Dissection revealed that migrating VIE pieces can accumulate in the internal organs, especially the kidneys. The clipped toe was successfully recognized only in 41% of checks. We conclude that photo-based identification is superior to both VIE-tagging and toe tipping in young agile frogs, and this method should be preferred in future studies seeking a cost-efficient yet reliable identification method in this species.

## 1. Introduction

Tracking individuals across time and space is key in research and conservation programs. There are various methods of individual identification, and choosing the right method for a particular goal is crucial for the success of the study. For example, the anatomical, physiological and behavioural features of a species may influence the suitability of a given identification method [1]. Different methods can significantly differ in costs in terms of money and time (e.g. expensive but easy-to-read microchips vs. inexpensive but time-consuming photography-based identification), hence limiting the options of projects on tight budget. Ethical considerations require to take into account the 3Rs principle (refinement, reduction and replacement) [2], and the ideal method should minimise adverse effects while maximising identification accuracy. Therefore, assessing the reliability of identification methods is important before starting long-term application in large numbers.

For many decades, toe clipping was the most conventional method of individual identification in amphibians. This is an inexpensive and fast method, but negative effects have been reported on various welfare-related endpoints such as stress hormone levels and behavioral changes as well as on long-term growth and survival [3], and clipped toes can reportedly regrow over time which limits identification success [4–6]. Clipping only the most distal phalanx (or just the toe pad in hylid frogs; both known as ‘toe tipping’) was proposed as an alternative to reduce the adverse effects, and this method has been used in different species [7,8]. Yet, studies on both the actual reduction of adverse effects and the reliability of toe tipping for individual identification are missing. Methods that are considered less invasive require the injection of extraneous material under the skin [1]. One such material is visible implant elastomer (VIE), a colourful silicone-based material that is usually considered harmless, with no effect on stress, movement or growth [9–16]. However, reports from several amphibian studies suggest that VIE may migrate under the skin, its colour may fade with time, and the material may even be rejected by the body, leading to identification failures [4,9,17–19]. Retainment of VIE tags may depend on the body part in which it was applied [4,18,20]. Studies that compared the reliability of toe clipping and VIE tagging in anurans came to contradicting conclusions [4–6,9], suggesting that the suitability of each method for individual identification depends on the species and circumstances of the study.

In contrast to the above methods, photo-based identification relies on the distribution of natural macroscopic colour patterns of the amphibian skin. Such patterns remain largely stable at least during adult life [21,22], thus providing reliable alternative to artificial tagging methods in various species [23–26]. By far, this method is the least invasive, and it requires only a suitable photographing device. However, when the number of individuals is more than a handful, photo-based identification is time-consuming and difficult to use on-the-spot, because the individual needs to be compared to all the previously taken photographs. Finding the matching images can be aided by various software, some of which are pricey though, and their suitability may greatly differ for a given set of images [24,25].

Reliability of different individual identification methods may greatly vary between adult and young life stages. Because juveniles have faster growth rates and potentially better regeneration capabilities, and skin coloration may change with ontogeny, methods that are reliable in adults may not necessarily work as well in young animals [27,28]. In this study, we assessed the reliability of three low-cost identification methods in young agile frogs (*Rana dalmatina*): toe tipping (clipping only the most distal phalanx), VIE tagging, and photo-based identification. We applied all three methods on recently metamorphosed froglets and followed the retention of individual markings for 488 days, including one winter hibernation period. We kept the frogs in small housing groups, where the application of only one VIE tag and one toe mark per individual was required for following the animals’ true identities. This enabled us to record mark retention by each method with high accuracy. Our study demonstrated that natural marks were far more stable than the applied artificial marks, and photo-based individual identification was always successful in young agile frogs, both when comparing small and large numbers of individuals.

## 2. Methods

### 2.1. Data collection

We used 113 agile frogs for this study, which we raised from the egg for the purposes of a long-term experiment. The eggs were collected on 15 March 2023 from 10 egg masses spawned naturally in our study pond (47.551195°N, 18.926682°E). The embryos and tadpoles were raised as described in [29]; after the third week of larval development until metamorphosis they were kept in mesocosms as described in [30], in groups of six siblings. When an animal reached the start of metamorphosis (appearance of forelimbs), we assigned an individual identifier (‘animal ID’) to it, photographed it in shallow water (either by digital camera or smart phone), and housed it in an individually-labelled plastic box until marking. After marking (see below), we housed the froglets in groups of five or six siblings outdoors, except for the winter during which they hibernated in a climate chamber [31]. Group sizes decreased with mortality during the experiment, and animals that died before marking or were excluded due to other reasons (S1 Appendix) were not counted in the N=113 for the present study. Details on housing conditions as well as animal exclusions from the study are described in S1 Appendix.

After all individuals had completed metamorphosis, we photographed them and marked them with both toe tipping and VIE tagging (Table 1). At tagging, the froglets had 0.6 – 1.3 g body mass and 19.6 – 26.4 mm snout-vent length (SVL; measured by Fiji version 1.54g [32] based on photographs). Each froglet was anesthetized by immersing in a 1 g/L water bath of tricaine-methanesulfonate (MS-222; a safe and humane anaesthetic widely used for amphibians [33–35]) buffered to neutral pH with the same amount of disodium hydrogen phosphate. When reflexes were lost (after ca. 2 minutes in the room-temperature solution), we placed the animal on wet paper towel for marking. We used flame-sterilized sharp scissors to remove one toe tip (the most distal phalanx) following a toe-clip coding system (Fig 1), such that every frog in a housing box would have a different toe tip removed. Additionally, we injected a VIE tag (Northwest Marine Technology, Inc., Shaw Island, WA, USA) of one out of six colours under the skin on one out of three body parts either on the left or the right side (Fig 1) following the manufactureŕs guidelines (but see S2 Appendix Fig S1 for signs of differential viscosity between the applied tags). Building on previous reports on varying tag retention [4,18,20], we chose body parts that are relatively far away from the lymph sinuses: the upper arm, the shin, and the foot at the edge of the sole. Another reason for choosing these three body parts was that they are relatively easy to see without picking up the animal, which might facilitate individual identification in the field, although in our study the tags were always examined with the animal being held in hand. We did not tag the toes or foot webs because these were too small. Length of the applied VIE tags varied between ca. 3 and 9 mm, and they were in general ca. twice as long in the shin than in the upper arm, because tag size was limited by the short length of the latter. Because VIE tags have been reported to move under the skin of frogs, we applied VIE in the same body part for all individuals within a housing box; therefore, the original body part remained unambiguously known for each individual, regardless of later tag relocations. However, the combination of tag colour and side (left/right) was different for all individuals housed in the same box. After marking, the animal was placed on its belly with legs unfolded to a loosely stretched posture for better visibility, and the full dorsal body surface was photographed along with its ‘animal ID’ using a Canon EOS R50 camera. The froglets were kept wet until they recovered from anaesthesia; all animals survived the procedures and woke up after ca. half an hour.

**Table 1.**
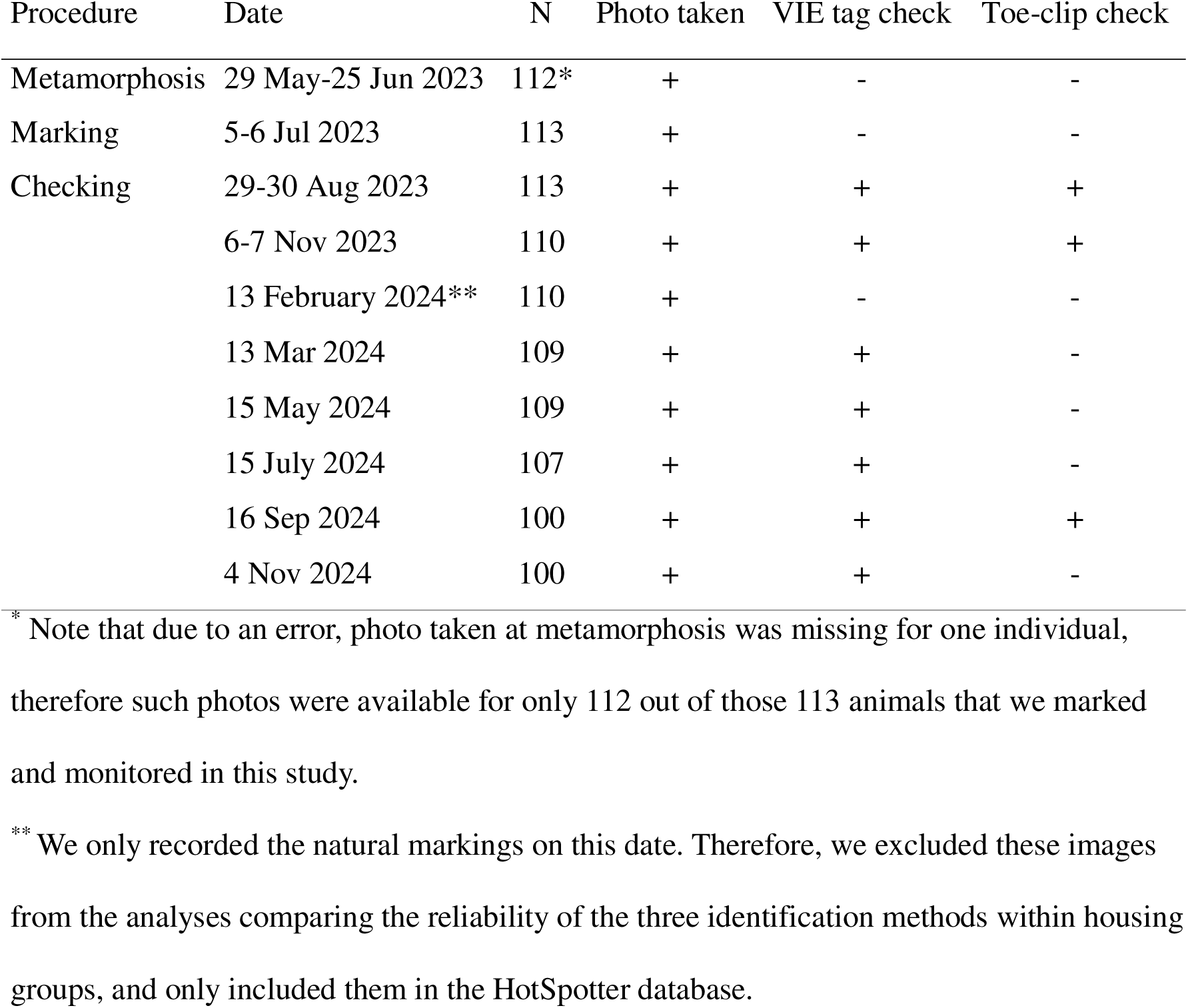
Timing of procedures and number of individuals (N) throughout the study.

**Fig 1.**
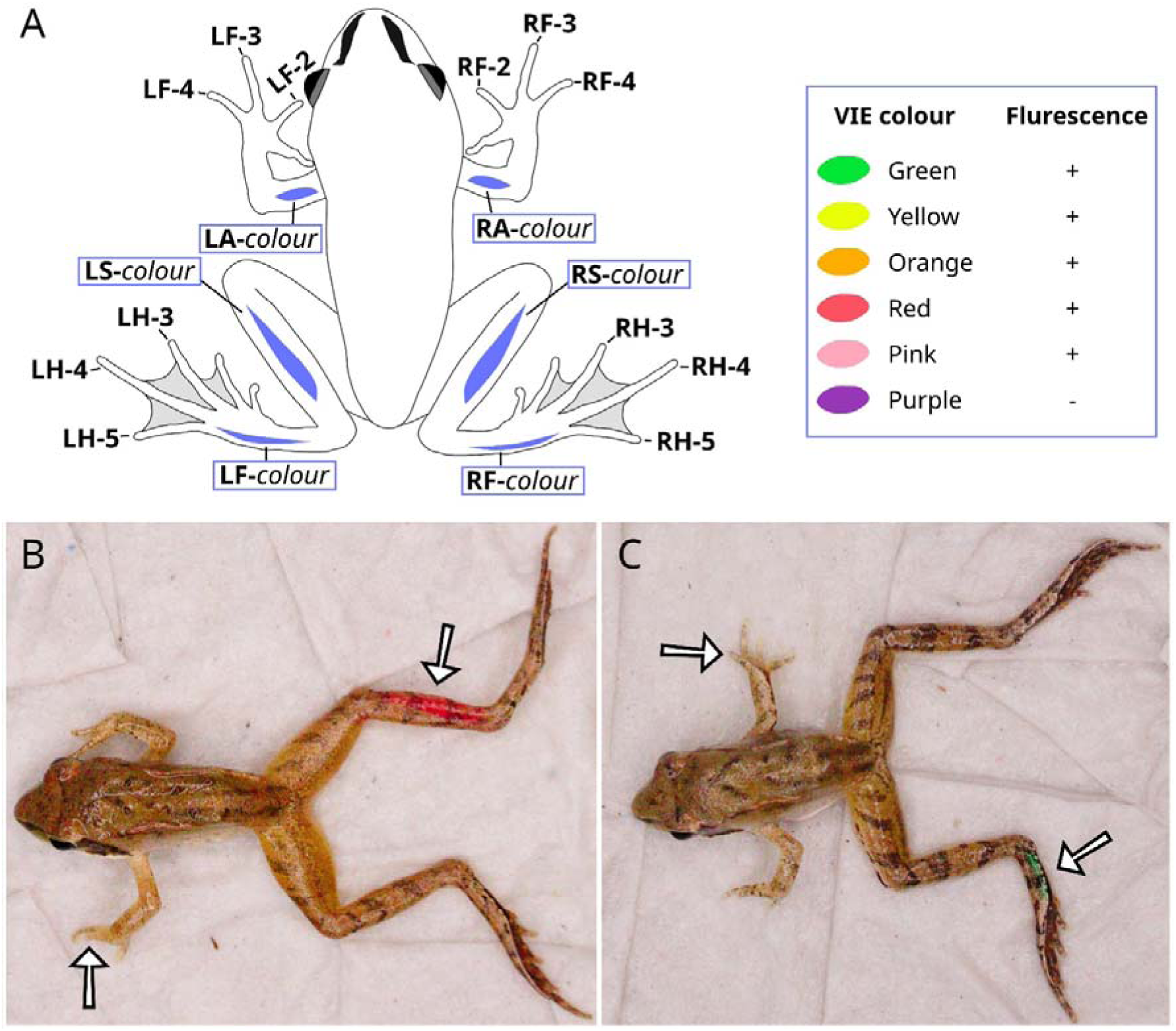
Individual identification by toe tipping and VIE tagging. Each individual was marked by the removal of one phalanx and the injection of one VIE colour into one of the highlighted body parts (A). VIE codes are highlighted with blue frame. Each marking code indicates the side of the body (L: left, R: right). Additionally, toe-clip codes indicate the limb (F: forelimb, H: hind limb) and digit (according to proximity to the body), while VIE codes indicate the body part (A: upper arm, S: shin, F: foot). The applied VIE colours and whether or not they are fluorescent under ultraviolet lighting are indicated on the right. Example photos show LF-4 combined with RS-red (B) and RF-4 combined with LF-green (C) just after tagging. Exposure curves were applied for each image to facilitate pattern visibility.

We evaluated the reliability of each identification method for over at least 14 months after marking, up until the start of the froglets’ second winter hibernation (Table 1). Checking events took place every two months, except for the duration of winter hibernation. During each checking event, we removed all frogs from each housing box in a haphazard order, and to each animal we assigned a number between 1 and the total number of frogs still alive (‘sampling ID’). Two researchers tried to identify the clipped toe and the colour and location of the VIE tag for each frog. They searched for VIE tags both with and without a UV-A torch light, because most of the VIE colours we used were fluorescent (Fig 1). At the first checking event (August 2023), researchers were allowed to discuss their decisions regarding VIE tags, while across the rest of the checking events, the two researchers independently assessed both VIE tags and toe-tip clips. They also recorded if their identifications of VIE colours and clipped toes were confident or uncertain. Lastly, they guessed the ‘animal ID’ for each froglet (‘on-site ID’) based on a combination of the available data, including the VIE tags, clipped toes, and exclusion (i.e. when markings were not detected on a single frog in the box, that frog could be identified as the only missing ‘animal ID’ in that box). Note that the sole purpose of the ‘on-site ID’ was to ensure that we could determine the true identity of each animal, that is essential for comparing mark retentions (see also the next paragraph). Because toe-tip clips were highly ambiguous during the first two checking events (see Results), after that we stopped evaluating mark retention by toe-tipping, except for the penultimate checking event when the froglets weighed 2.7 – 7.5 g with 30.6 – 44.5 mm SVL (Table 1).

At each checking event, the froglets were photographed together with their ‘sampling ID’ while positioning themselves freely in clean water [26] either by a Sony Cyber-shot DSC-HX200V under varying light conditions indoors, or by a Canon EOS R50 digital camera equipped with circular polar filter in a custom-developed photo box (light: 5000K Matcheasy LED strip, 5V, 3m). Later, we brightened the images by applying exposure curves in RawTherapee (version 5.9; https://rawtherapee.com) to enhance the visibility of skin patterns. Then, an experienced researcher who was blind to the ‘on-site ID’ compared the images of frogs in a housing box to the photographs captured at marking of all froglets housed in the same box, and assigned a ‘photo-based ID’ to each individual at each checking event. Each photo was compared with all individuals photographed at marking from the same housing box (regardless of being identified already, and being dead or alive), so exclusion was never used during photo-identification. VIE tags were rarely visible on the images taken during the checking events; for such cases, the researcher was instructed to ignore VIE tags during photo-identification and only consider the skin patterns of the frogs.

To assess the agreement of photo-identification within the small housing groups between different observers, a subset of the images was evaluated by a second, less experienced researcher. For this purpose, the 20 housing boxes were randomly distributed across the seven checking events, such that a similar number of frogs would be chosen from each event, and each of the 113 frogs would be chosen only once. Additionally, to investigate whether the photographs allow individual identification already at the start of metamorphosis, these photographs were compared to the images captured in August 2023; this was done by the experienced researcher for all frogs (N = 113), and by the less experienced researcher for a subset of two frogs picked randomly from each housing box (N = 40). For all photo-identifications, the observers also recorded if they were confident or uncertain. For the photos taken at metamorphosis, the experienced researcher scored picture quality as good, medium, or poor (Fig S2), based on the visibility of skin markings due to both image properties (e.g. sometimes the animal was not in focus or partially hidden by reflections on the water) and the posture of the animal (e.g. the tail sometimes partially covered the legs). These quality issues occurred because, for logistical feasibility, the metamorphs were photographed by several researchers, none of whom had experience with capturing high-quality pictures for individual identification. All photos taken after marking had good quality, with all skin markings being fully visible, as all these pictures were taken by a single, experienced researcher.

To evaluate if the natural markings enabled individual identification in a large group of juvenile frogs as well (and not just among five or six animals as above), we performed computer-assisted identification in HotSpotter [36] using images from two different dates per individual. To avoid disturbing effects of the background [26], we used images from those two occasions when the animals were photographed in water against a uniform white background: November 2023 and February 2024. First, we added one photo of each individual from November and assigned the ‘animal ID’ to each image in the ‘name’ column in HotSpotter. Then, we added a photo of each animal from February without assigning any ID to it. To maximise the size of this dataset, we used images from all frogs still alive (N=112 in November 2023 and N=111 in February 2024, including two animals that were excluded from all other analyses due to having two VIE tags). First, we carried out a ‘software-performance test’ by running ‘queries’ in HotSpotter for the 111 individuals photographed in February, when we recorded the rank of the true match from November for each queried image. Meanwhile, the author who created the database checked if VIE tags were visible on the included images, and listed those individuals whose images from neither November nor February featured visible VIE (N=91). Finally, we performed an ‘observer test’ focusing on the latter 91 animals to assess if human observers could identify the matching images solely based on the natural colour markings of the animals. In the ‘observer test’, images from February were haphazardly assigned to one of two authors who had no previous experience in photo-based identification of these frogs (45 images queried by one and 46 queried by the other author). Their task was to assign the ‘animal ID’ to each queried image by evaluating the first 20 potential matches listed by HotSpotter (i.e. rank ≤ 20; a threshold also used in previous studies [26, 27, 37]).

Mortality post-marking was low (N = 11) and happened during the warmest times (Table 1; note that two further individuals were taken out of the study after 5 and 6 checking events, respectively, for reasons not related to this study). The froglets that died after marking were dissected and examined for VIE material in their internal organs using an APOMIC SHD200 digital microscope.

### 2.2. Data analysis

First, we investigated which method yielded the most unambiguous decisions regardless of the correctness of identification. To this end, we categorized each identification attempt as ‘unambiguous’ in cases where only a single, confident guess was made, and ‘ambiguous’ where the observers were uncertain or in conflict with each other, or they could not find the marking at all. We used a generalized linear mixed-effects model with binomial error and logit link, where the dependent variable was the binary ambiguity category. The fixed factor was the type of method (toe-clip, VIE tag, photo), and the random factor was ‘animal ID’ (inferred from ‘photo-based ID’ as explained below). Because this analysis showed that both VIE and toe-tip markings were often ambiguous (see Results), we could not use the ‘on-site ID’ as the “true” individual identifier. Therefore, we used the ‘photo-based ID’ as the “true” identification of ‘animal ID’ in the second set of analyses, considering the following reasons. During photo-identification (using the images taken at marking as reference), the observers were never in conflict and always found a putative match, which always matched the ID assigned based on the only tag colour that was always unambiguously recognizable when found (see Results). In each case where the ‘on-site ID’ did not match the ‘photo-based ID’, we double-checked if there was any error made during photo-identification; in all these cases the photo belonging to the non-matching ‘on-site ID’ clearly showed a different skin pattern compared to the image taken at marking, meaning that the ‘photo-based ID’ was correct and the ‘on-site ID’ (based on VIE and toe-tip clips) was mistaken.

To analyse the success rate of identification by toe-tip clips and VIE tags, we used the same model structure as above, but the binary categories of the dependent variable were ‘successful’ (i.e. the assigned ID was the same as the ‘photo-based ID’) versus ‘failed’ (i.e. the assigned ID was conflicted, or not found, or did not match the ‘photo-based ID’). For toe-tip clips, the fixed effects were the digit clipped (regardless of side) and the checking event (we used the latter as categorical variable to allow for identification success to change non-linearly over time). For VIE tags, the fixed effects were tag colour, the tagged body part (regardless of side), and checking event. We calculated the identification success by VIE tags in three alternative ways. First, we considered all identifications successful when the colour of the tag was correctly recognized, regardless of tag location. Second, we considered all identifications successful when the location of the tag was correct regardless of tag colour; tag location was recorded as correct when it was found in the body part where it had been injected into, and incorrect if it was not found at all or found in other part(s) of the body (i.e. further than the nearest joints from the initial location). Third, we considered all identifications successful when both the colour and location of the tag was correct (i.e. the right colour was found in the body part where it had been injected).

For all statistical analyses, we used the R 4.3.2 software, with function ‘glmer’ of package ‘lme4’ for binomial models. We used the ‘DHARMa’ package for residual diagnostics to check that the statistical requirements of our models were met. From each model, we calculated post-hoc pairwise comparisons using the ‘emmeans’ package, correcting the P-values for multiple testing with the false-discovery rate method [38].

### 2.3. Ethics statement

This study was approved by the Ethics Committee of the Plant Protection Institute and licensed by the Environment Protection and Nature Conservation Department of the Pest County Bureau of the Hungarian Government (PE-06/KTF/00754-8/2022, PE-06/KTF/00754-9/2022, PE-06/KTF/00754-10/2022, PE/EA/295-7/2018, PE/EA/00270-6/2023).

## 3. Results

Ambiguity rate differed significantly between each of the three methods: it was highest for toe tipping and lowest for photo-identification (Table 2; P < 0.0001 for all pairwise differences). The nature of ambiguity also varied by method (Table 2). For toe-tip clips, the two observers often made conflicting identifications or recorded their guesses as uncertain (Table 2). For VIE tags, these kinds of ambiguity were less frequent, but sometimes the tags were not found at all (Table 2). For photo-identification based on the images taken at marking, ambiguity was rare and always occurred in the form of the observers being somewhat uncertain about their choice (Table 2). When the images taken at metamorphosis were used as the basis for photo-identification, the observers were often uncertain and sometimes found conflicting or no matches at all (Table 2). The dataset containing decisions for each individual at each checking event is available on Figshare [39].

**Table 2.**
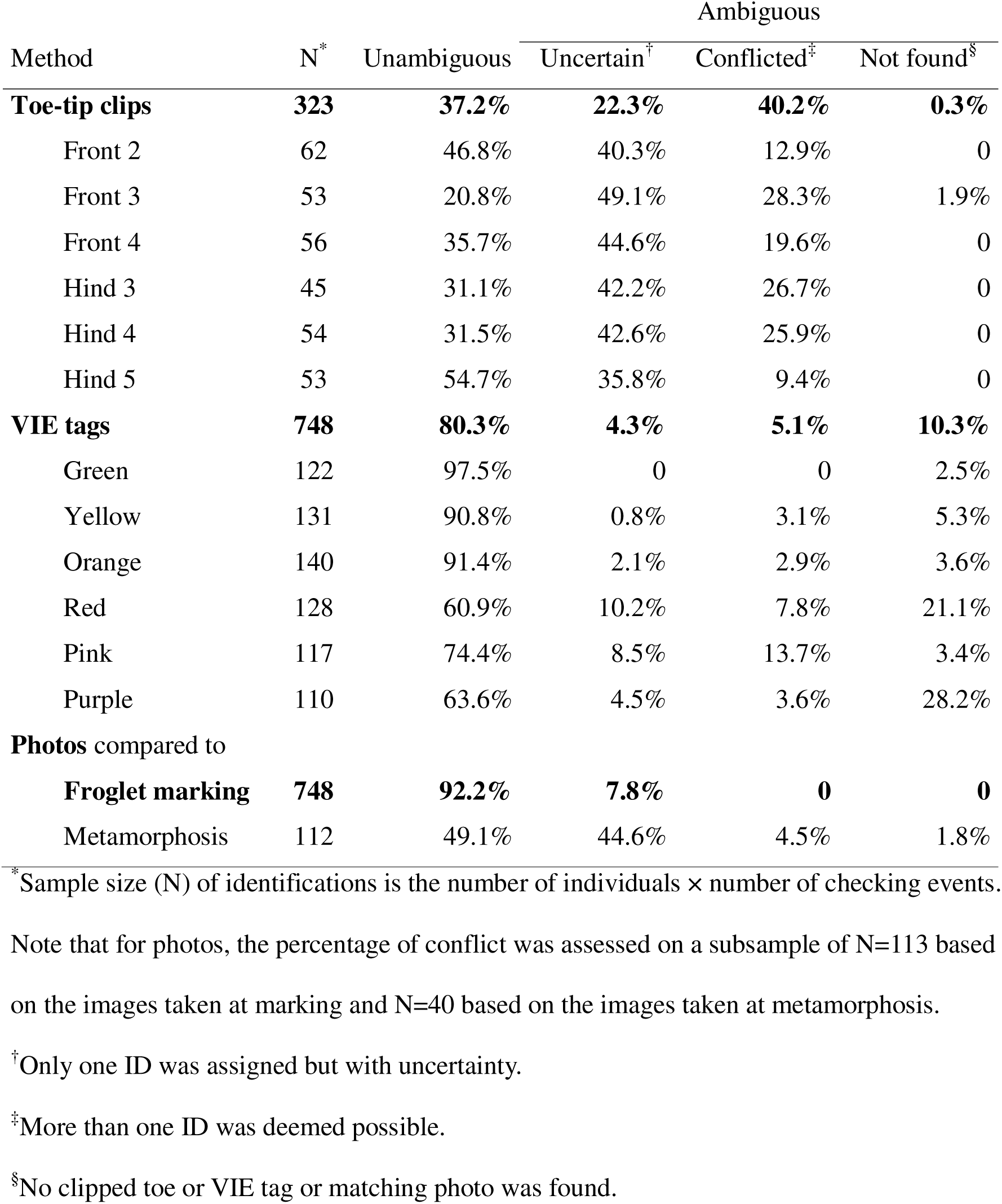
Percentage of ambiguous and unambiguous identifications for each marking method, regardless of correctness of the identification.

When the VIE tag was found, green was the only colour that was always recognized without ambiguity (Table 2). For the green-tagged individuals, every ‘photo-based ID’ matched the guess based on the VIE tag (N = 119). For other tag colours, whenever there was a mismatch between the ‘photo-based ID’ and the guess based on the VIE tag (N = 92 out of 552 individual checks where the tag was found), the photograph taken at marking always matched the individual assigned by ‘photo-based ID’ and not the individual assigned by VIE tag. Because of these reasons, we treated all identifications based on the images taken at marking as correct, and we defined the success of the other methods based on the ‘photo-based ID’.

The overall success of identification by toe tipping was only 40.9%, varying from 15.1% to 67.9% across digits (Fig 2) and from 35% during the first two checking events to 54% in September before the second hibernation (Fig 2). For VIE tags, identification based on colour was correct in 77.4% of cases, with significantly higher success for green, yellow, and orange than for red, pink and purple tags (Fig 3). However, out of the 671 individual checks where the tag was found, in 102 cases it was not located in the body part where it had been injected (15.2%). In 77 cases (10.3%) involving 36 individuals the tag was not found at all, but in 16 of those individuals it was found again 1-4 checking events later (including 8 individuals in which the tag disappeared and was found again more than once). Dissection of the froglets that died or were taken out of the study revealed that, in 5 out of 13 animals, some of the VIE material moved into the internal organs (Fig S3); this happened from all three body parts where the tag had been injected. We recorded tag dislocation including complete disappearance in 179 out of 748 individual checks (23.9%), and it was significantly more frequent in the arms than in the legs (Fig 3), for the purple and red tags than for other colours (Fig 3), and at later checking events (Fig 4). Altogether, when both the colour and the location of the VIE tags were considered, identification success was 65.4% overall (Fig 3), decreasing from 78.8% to 51% from the first to the last checking event (Fig 4).

**Fig 2.**
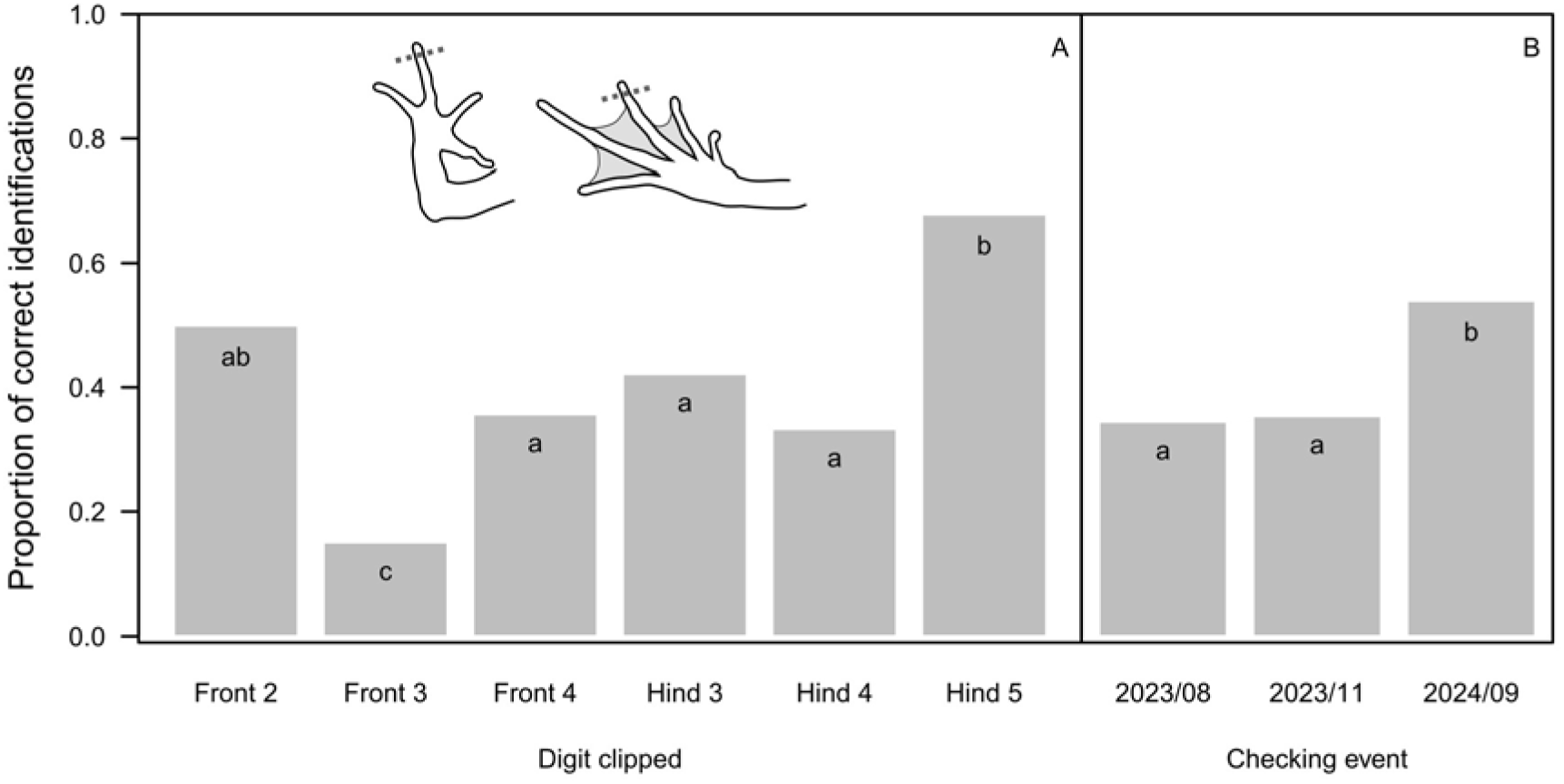
Identification success for marking by toe tipping, depending on the digit clipped (A) and the timing of checking the markings (B). Letters at the top of bar plots indicate statistically significant differences (groups with different letters differ at P < 0.05 after correcting for false-discovery rate).

**Fig 3.**
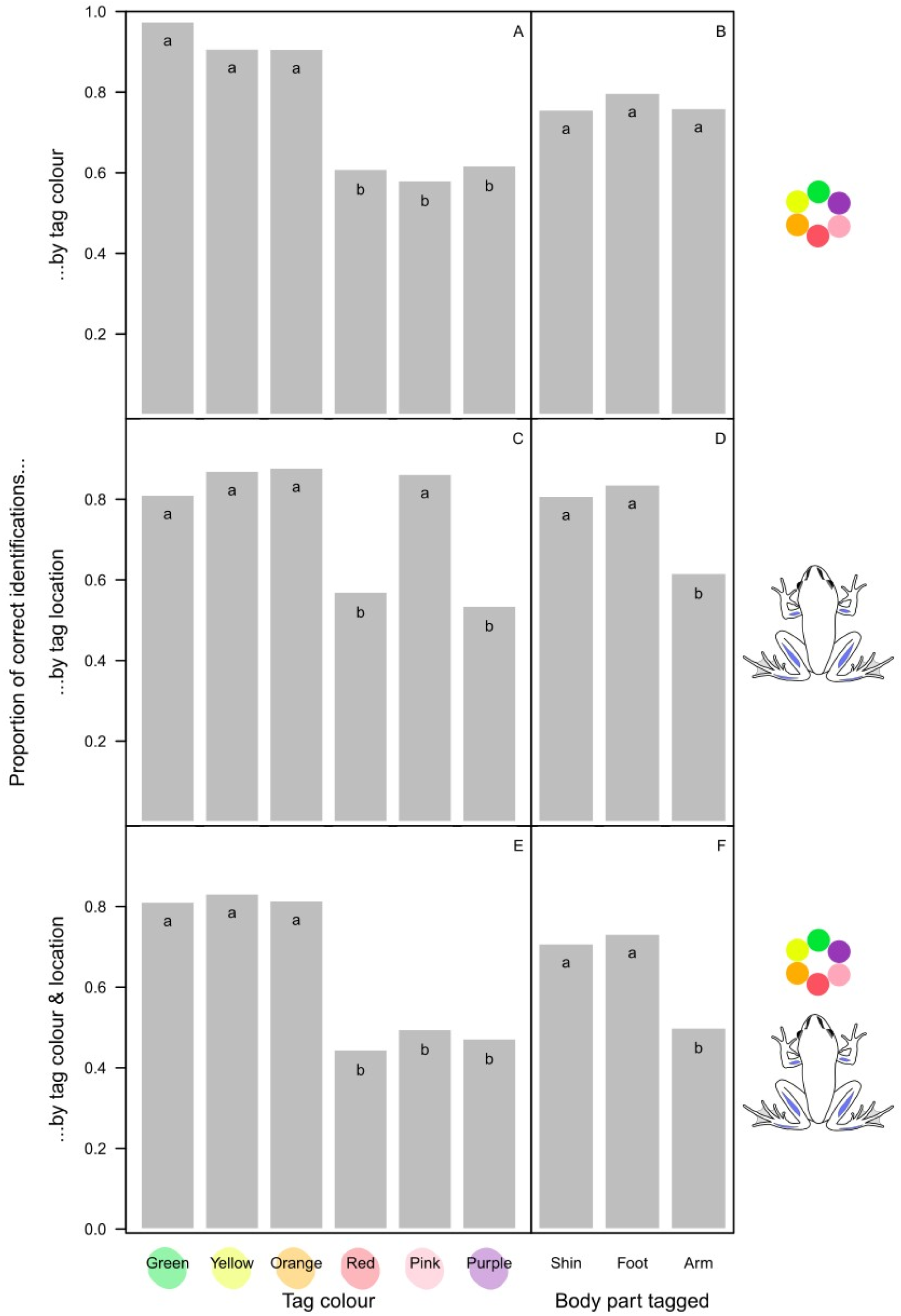
Identification success calculated in three different ways for marking by VIE tagging, depending on tag colour (panels A, C and E) and tagged body part (panels B, D and F). Letters at the top of bar plots indicate statistically significant differences (groups with different letters differ at P < 0.01 after correcting for false-discovery rate).

**Fig 4.**
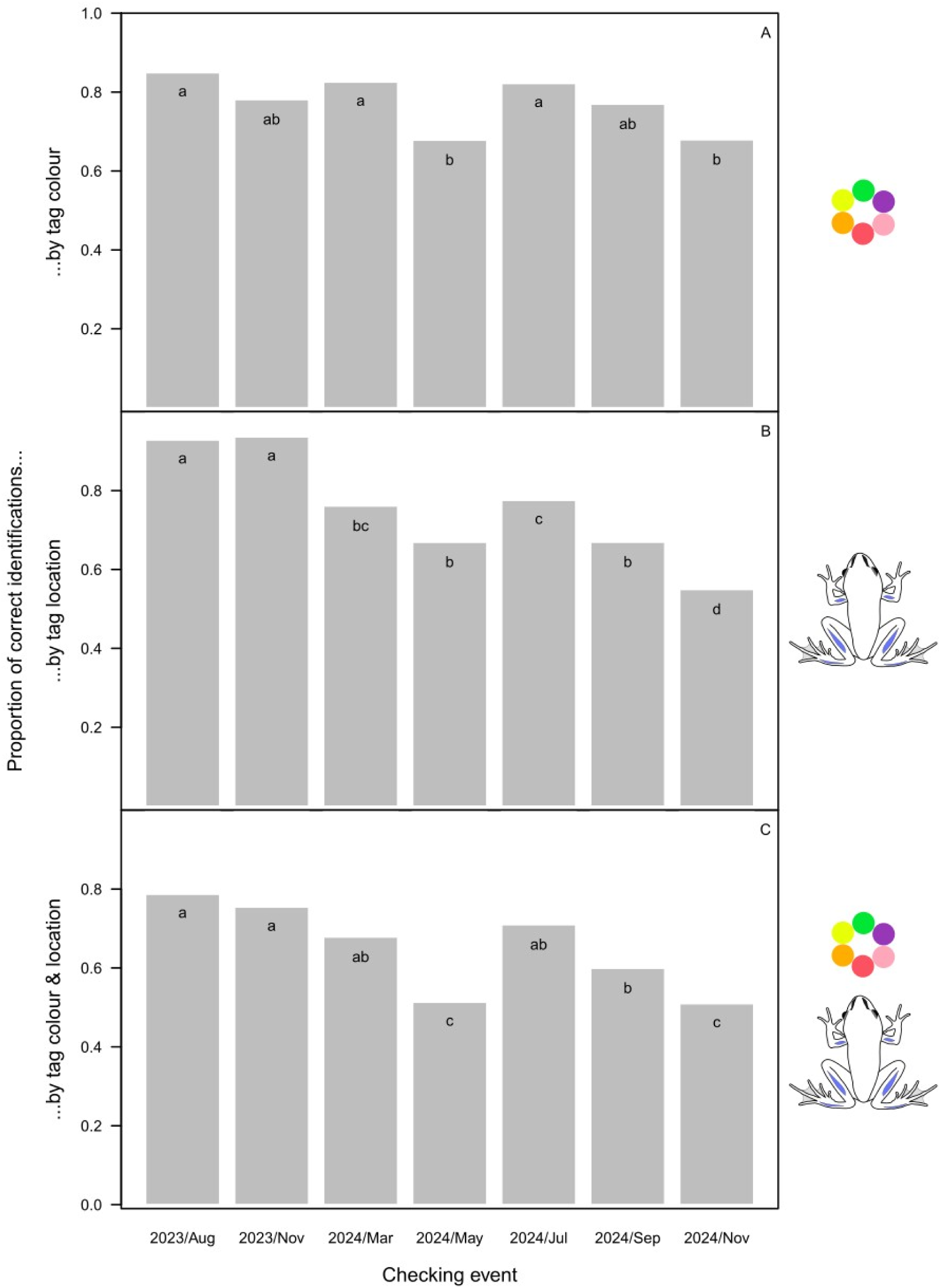
dentification success calculated in three different ways for marking by VIE tagging, depending on the timing of checking the markings. Letters at the top of bar plots indicate statistically significant differences (groups with different letters differ at P < 0.01 after correcting for false-discovery rate).

When comparing each of 111 queried photos from February 2024 with all the other images in the ‘software-performance test’, the image taken of the same animal three months earlier was identified as the most likely match (rank=1) by HotSpotter for 75.7% of the queried images, 92.8% received a rank ≤10, and 98.2% received a rank ≤ 20. Considering only the results of those 91 individuals whose photos did not feature any signs of VIE, 73.6% received rank=1, 92.3% received rank ≤ 10, and 98.9% received rank ≤ 20; the only individual with rank > 20 received rank = 48. In the ‘observer test’, the ‘animal ID’ was correctly assigned to each of the 90 individuals with rank ≤ 20, and the observer dealing with the one individual whose matching pair was not present among the rank ≤ 20 images correctly recognized that there was no ‘animal ID’ to assign. With other words, the overall false rejection rate was less than 2% (due to software performance), with a false acceptance rate of 0% by the observers. The dataset containing all software-performance and observer results, and screenshots from HotSpotter are available on Figshare [39].

Based on the photographs captured at metamorphosis, individual identification was highly successful (93.8%) within the small groups of co-housed animals despite the relatively high uncertainty reported by the observers. However, 47.3% of the photos taken at metamorphosis had some problems with image quality or the body posture of the animal or both, and 42.0% had poor quality (i.e. the problems resulted in low visibility of the leg patterns). Ambiguity rate decreased as photo quality improved from poor (70.2%) through medium (37.7%) to good (33.3%; χ^2^ test: χ^2^ = 12.17, df = 2, P = 0.002). Identification success was 91.5% for poor-quality photos, 94.3% for medium-quality photos, and 100% for good-quality photos. The dataset containing image qualities and identification results of metamorphs is available on Figshare [39].

## 3. Discussion

In our study, photographs of the natural pigment markings on the skin were clearly superior to both toe tipping and VIE tagging for individual identification of agile frogs during the first one and a half years of their post-metamorphic life. When image quality was good, the observers had low ambiguity when assessing potential matches, and the matches they assigned were always correct when we were able to assess this with a reliable alternative marking (i.e. green tags that were unambiguously detected in our study). Although the skin patterns sometimes changed slightly as the animals aged (S2 Appendix Fig S4), the observers were always able to find the correct match from up to six candidate individuals in our study system. Reliability of computer-assisted identification in the large dataset (92% with rank ≤ 10 and 99% with rank ≤ 20) was comparable to previous reports on successful photo-based identification in amphibians [24–27, 39]. This result, together with the fact that all four observers (two focusing on the small groups and two on the HotSpotter database) always identified the matching photos if those were present among the list of potential matches that they evaluated, demonstrates that natural colour markings of agile frogs are unique already in their first year of life. Substantially different body postures on matching images likely constrained the performance of HotSpotter in our study. By adding multiple images per individual [25,40], and applying optimized techniques for image capturing while paying more attention to animal positioning and lighting conditions [26], performance of computer-assisted photo-identification is expected to improve even further. Indeed, using the latter approach, we found in a parallel study on adult agile frogs that 100% accuracy can be achieved with computer-assisted photo-identification in datasets consisting of several hundred individuals [26]. Furthermore, in the present study, the photographs taken at metamorphosis (i.e. emergence of front limbs) also proved suitable for individual identification of the young frogs 2-3 months later, when photos of ≤ 6 co-housed siblings were compared. When picture quality was good, photo-based identification was 100% successful regardless of the timing of taking the reference photographs (i.e. at metamorphosis or at froglet marking), and it was fairly high even with the poorest-quality pictures taken at metamorphosis (91.5%). This suggests that the unique individual patterns of agile frogs are already recognizable at very early life stages and remain largely stable throughout and after metamorphosis (S2 Appendix Fig S5), similarly to what has been found in a few other anuran species [41,42]. The drawback of the photo-identification method is that comparison of the individual to the reference photographs requires the availability of those photographs (e.g. on a computer at hand) and non-negligible amount of time that increases with the number of potential matches that need to be checked. For example, in our study, on-site assessment of VIE tags and toe-tip clips took qualitatively similar time to photographing, but the latter had to be followed by off-site image processing and subsequent photo comparison. Nevertheless, the time needed for finding the right match can be sped up by computer-assisted photo-identification [43], using suitable software [24,25,40] and optimized photography methods [23,26]. Earlier studies that used computer-assisted photo-identification in amphibians had widely variable success rates (Table S1 in [26]), which might be due to variability in the photography settings and software applied or the stability of skin markings.

Overall, the second-best method in our study was VIE tagging. In our system, tag colour alone should have been enough for successful individual identification because the same colour was rarely injected into more than one animal in the same box, and the tags rarely ever moved from one side of the body to the other. However, we had difficulty with telling apart three of the six colours (red, pink, and purple), probably because the skin of the tagged body parts is reddish-brown (S2 Appendix Figs S4 and S5) and sometimes can be quite dark. In this respect, body parts with lightly coloured skin would be preferable for VIE tagging; however, the only such area on agile frogs is the belly and the ventral side of the thigh, where tag retention has been typically low in other species [18,20]. Furthermore, sometimes we did not even find the tag, despite that we were often able to locate the same tags during some of the subsequent checking events. The reason for the latter is that the tags often broke into several pieces, and the pieces migrated away from the injection point both towards and away from the centre of the body, often leaving only small, easy-to-miss dots close to the skin surface. These findings are similar to those reported from studies on other amphibian species [18,19,44]. Tag dislocation may further reduce success in situations where tag location is part of the individual identification code; for example, we found that tags injected into the shin can move into the foot and *vice versa*. In our study, dislocation was more frequent for tags injected into the upper arm compared to the shin and foot; however, this contrasts findings in other anuran species [20,28,45] and might be explained by tag size rather than by location, because we injected smaller tags into the upper arms due to their smaller size relative to the two target sites on the legs. However, larger tags were more likely to migrate in other amphibians [46]. Altogether, in our study, tag migration resulted in the 77% identification success based solely on tag colour dropping to 65% based on the combination of tag colour and location. This overall low success rate would be unacceptable for many studies; however, VIE tags could be still reliable for applications that do not require a large number of individual codes. For example, our data suggest that young agile frogs could be successfully batch-marked with green, yellow, or orange VIE for tracking cohorts.

Our data on tag migration raise concerns about the health risks associated with VIE tagging. Regardless of initial tag location and colour, some of the VIE material moved into the internal organs, mostly the kidneys, which agrees with findings on adult cane toads (*Rhinella marina*; [47]). In the latter study, VIE in the kidney was associated with inflammation and damage to the vasculature that progressed in severity over time. This is especially worrisome in light of growing concerns about the ecotoxicological effects of microplastics [48]. Notably, although growth and survival were reported to be unaffected by VIE tagging in various amphibians, this has been studied over relatively short periods, rarely longer than a year [9–15]. Therefore, caution is recommended with using VIE especially in free-living wild animals, and further research is needed to validate its long-term safety.

To our knowledge, this is the first report on mark retention by toe tipping under controlled experimental conditions, despite its reported usage in the field [7,8]. Removal of only the most distal phalanx yielded unacceptably low identification success in our study, ranging between 15.1% (front 3) and 67.9% (hind 5) depending on the marked toe. This may be due to particularly high regeneration capacities in young juveniles, although 100% toe-clip retention was observed in boreal chorus frogs (*Psuedacris maculata*) over one year after metamorphosis [9]. However, the length of the clipped part was not clarified in the latter study; studies that reported high reliability of toe-clipping typically removed more than one phalanx per toe (e.g. [4,44]) whereas we removed only the most distal phalanx. Nevertheless, even if identification success would be increased by more extensive clipping, negative consequences of toe-clipping on survival can increase with the severity of mutilation (e.g. by the number of toes clipped per animal, which inevitably increases when a large number of individuals have to be marked uniquely; [9,49, 50]). Our results suggest that the practice of clipping just one phalanx per toe may lead to the data turning out useless, and the mutilation having been done in vain. As pointed out by a recent systematic review on the welfare impacts of toe-clipping amphibians [3], data to help decisions in this ethically loaded topic are still too scarce and heterogeneous. While the debate is still going on and more data are being collected, we recommend using less invasive methods for marking individuals whenever feasible.

In conclusion, our study shows that photo-based individual identification, besides serving refinement according the 3Rs [2], is superior to both VIE-tagging and toe tipping for individual identification in young agile frogs. When the images were taken with care to ensure good quality, photo-identification had 100% success over one and a half years in the housing groups, and major individually unique patterns were recognizable already at metamorphosis. Although our animals were kept in small groups, the high stability of skin patterns and the success of the computer-assisted identification for 91 froglets suggest that similarly exquisite performance may be achieved with larger numbers of individuals. In contrast, both VIE-tagging and toe tipping showed lower identification success, and the emergence of VIE in internal organs suggests health risks. We encourage the usage of the most reliable and least invasive, even though arguably most time-consuming photo-based method in future studies on agile frogs whenever possible.

## Acknowledgements

We thank Árpád Ferincz for the VIE material, and Emese Balogh, Judit Baumann, Zoltán Örkényi, Mihály B. Rusz and all colleagues and students at the Department of Evolutionary Ecology for their help during the study.

## Supporting Information

**S1 Appendix. Details of rearing conditions and exclusions from the experiment.**

**S2 Appendix. Supplementary figures.**

## References

1. Murray DL, Fuller MR. A critical review of the effects of marking on the biology of vertebrates. In: Boitani L, Fuller TK, editors. Research Techniques in Animal Ecology: Controversies and Consequences. New York: Columbia University Press; 2000. pp. 15–64.

2. Russell WMS, Burch RL. The principles of humane experimental technique. UFAW, Wheathampstead, UK: Universities Federation for Animal Welfare; 1992. Available: https://caat.jhsph.edu/the-principles-of-humane-experimental-technique/

3. Zemanova MA, Martín RL, Leenaars CHC. The impact of toe-clipping on animal welfare in amphibians: A systematic review. Glob Ecol Conserv. 2025;59: e03582. doi:10.1016/j.gecco.2025.e03582

4. Hoffmann K, McGarrity ME, Johnson SA. Technology meets tradition: A combined VIE-C technique for individually marking anurans. Appl Herpetol. 2008;5: 265–280. doi:10.1163/157075408785911002

5. Campbell TS, Irvin P, Campbell KR, Hoffmann K, Dykes ME, Harding AJ, et al. Evaluation of a new technique for marking anurans. Appl Herpetol. 2009;6: 247–256. doi:10.1163/157075409X420042

6. Davis TM, Ovaska K. Individual recognition of amphibians: effects of toe clipping and fluorescent tagging on the salamander *Plethodon vehiculum*. J Herp. 2001;35: 217–225.

7. Phillott AD, Skerratt LF, McDonald KR, Lemckert FL, Hines HB, Clarke JM, et al. Toe-clipping as an acceptable method of identifying individual anurans in mark recapture studies. Herpetol Rev. 2007;38: 305–308.

8. Kenyon N, Phillott AD, Alford RA. Evaluation of the photographic identification method (PIM) as a tool to identify adult *Litoria genimaculata* (Anura: Hylidae). Herpetol Conserv Biol. 2009;4: 403–410.

9. Swanson JE, Bailey LL, Muths E, Chris Funk W. Factors influencing survival and mark retention in postmetamorphic boreal chorus frogs. Copeia. 2013; 670–675. doi:10.1643/CH-12-129

10. Sapsford SJ, Roznik EA, Alford RA, Schwarzkopf L. Visible implant elastomer marking does not affect short-term movements or survival rates of the treefrog litoria rheocola. Herpetologica. 2014;70: 23–33. doi:10.1655/HERPETOLOGICA-D-13-0004

11. Fouilloux CA, Garcia-Costoya G, Rojas B. Visible implant elastomer (VIE) success in early larval stages of a tropical amphibian species. PeerJ. 2020;8: e9630. doi:10.7717/peerj.9630

12. Knapp DD, Diaz L, Unger S, Anderson CN, Spear SF, Williams LA, et al. Long-term retention, readability, and health effects of visible implant elastomer (VIE) and visible implant alpha (VI Alpha) tags in larval eastern hellbenders (*Cryptobranchus alleganiensis alleganiensis*). J Herpetol. 2023;57: 133–141. doi:10.1670/22-011

13. Bainbridge L, Stockwell M, Valdez J, Klop-toker K, Clulow S, Clulow J, et al. Tagging tadpoles: retention rates and impacts of visible implant elastomer (VIE) tags from the larval to adult amphibian stages. Herpetol J. 2015;25: 133–140.

14. Phillips CT, Fries JN. An Evaluation of Visible Implant Elastomer for Marking the Federally Listed Fountain Darter and the San Marcos Salamander. North Am J Fish Manag. 2009;29: 529–532. doi:10.1577/m08-138.1

15. Oropeza-Sánchez MT, Sandoval-Comte A, García-Bañuelos P, Hernández-López P, Pineda E. Use of visible implant elastomer and its effect on the survival of an endangered minute salamander. Anim Biodivers Conserv. 2020;43: 187–190. doi:10.32800/abc.2020.43.0187

16. Narayan EJ, Gramapurohit NP. Urinary corticosterone metabolite responses to capture and visual elastomer tagging in the Asian toad (*Duttaphrynus melanostictus*). Herpetol J. 2019;29: 179–183. doi:10.33256/hj29.3.179183

17. Schmidt K, Schwarzkopf L. Visible implant elastomer tagging and toe-clipping: Effects of marking on locomotor performance of frogs and skinks. Herpetol J. 2010;20: 99–105.

18. Brannelly LA, Chatfield MWH, Richards-Zawacki CL. Visual implant elastomer (VIE) tags are an unreliable method of identification in adult anurans. Herpetol J. 2013;23: 125–129.

19. Heemeyer JL, Homyack JA, Haas CA. Retention and readability of visible implant elastomer marks in eastern red-backed salamanders (*Plethodon cinereus*). Herpetol Rev. 2007;38: 425–428.

20. Moosman DL, Moosman Jr. PR. Subcutaneous movements of visible implant elastomers in wood frogs (*Rana sylvatica*). Herpetol Rev. 2006;37: 300–301.

21. Rojas B, Lawrence JP, Márquez R. Amphibian Coloration: Proximate Mechanisms, Function, and Evolution. In: Moreno-Rueda G, Comas M, editors. Evolutionary Ecology of Amphibians. Boca Raton: CRC Press; 2023. pp. 219–258. doi:10.1201/9781003093312-12

22. Price AC, Weadick CJ, Shim J, Rodd FH. Pigments, patterns, and fish behavior. Zebrafish. 2008;5: 297–307. doi:10.1089/zeb.2008.0551

23. Bendik NF, Morrison TA, Gluesenkamp AG, Sanders MS, O’Donnell LJ. Computer-assisted photo identification outperforms Visible Implant Elastomers in an endangered salamander, Eurycea tonkawae. PLoS One. 2013;8: e59424. doi:10.1371/journal.pone.0059424

24. Burgstaller S, Gollmann G, Landler L. The green toad example: A comparison of pattern recognition software. North West J Zool. 2021;17: 96–99.

25. Matthé M, Sannolo M, Winiarski K, Spitzen - van der Sluijs A, Goedbloed D, Steinfartz S, et al. Comparison of photo-matching algorithms commonly used for photographic capture–recapture studies. Ecol Evol. 2017;7: 5861–5872. doi:10.1002/ece3.3140

26. Nemesházi E, Mikó Z, Ujhegyi N, Kásler A, Lehofer N, Bókony V. Put the frog in water: simple methods for improving individual identification as demonstrated with the agile frog. Prepr BioRxiv. 2025. doi:10.1101/2025.02.25.640084

27. Aevarsson U, Graves A, Carter KC, Doherty-Bone TM, Kane D, Servini F, et al. Individual identification of the lake Oku clawed frog (*Xenopus longipes*) using a photographic identification technique. Herpetol Conserv Biol. 2022;17: 67–75.

28. Holcomb J, Lawrence K, Bailey S, Dittmer D. Considerations for and use of visible implant elastomers to track and estimate abundance of juvenile boreal toads (Anaxyrus boreas boreas) in Utah, USA. Herpetlogical Rev. 2022;53: 579–585.

29. Bókony V, Balogh E, Mikó Z, Kásler A, Örkényi Z, Ujhegyi N. Higher sex-reversal rate of urban frogs in a common-garden experiment suggests adaptive microevolution. Evol Appl. 2025;18: e70093. doi:10.1111/eva.70093

30. Bókony V, Üveges B, Móricz ÁM, Hettyey A. Competition induces increased toxin production in toad larvae without allelopathic effects on heterospecific tadpoles. Funct Ecol. 2018;32: 667–675. doi:10.1111/1365-2435.12994

31. Kásler A, Holly D, Herczeg D, Ujszegi J, Hettyey A. Chytridiomycosis and climate change: exposure to *Batrachochytrium dendrobatidis* and mild winter conditions do not increase mortality in juvenile agile frogs during hibernation. Anim Conserv. 2023;26: 654–662. doi:10.1111/acv.12851

32. Schindelin J, Arganda-Carreras I, Frise E, Kaynig V, Longair M, Pietzsch T, et al. Fiji: an open-source platform for biological-image analysis. Nat Methods. 2012;9: 676–682. doi:10.1038/nmeth.2019

33. Mitchell MA. Anesthetic considerations for amphibians. J Exot Pet Med. 2009;18: 40–49. doi:10.1053/j.jepm.2008.11.006

34. Downes H. Tricaine anesthesia in Amphibia: a review. Bull Assoc Reptil Amphib Vet. 1995;5: 11–16. doi:10.5818/1076-3139.5.2.11

35. Hadfield CA, Whitaker BR. Amphibian emergency medicine and care. Semin Avian Exot Pet Med. 2005;14: 79–89. doi:10.1053/j.saep.2005.04.003

36. Crall JP, Stewart C V., Berger-Wolf TY, Rubenstein DI, Sundaresan SR. HotSpotter-Patterned species instance recognition. Proc IEEE Work Appl Comput Vis. 2013; 230–237. doi:10.1109/WACV.2013.6475023

37. Morrison TA, Keinath D, Estes-Zumpf W, Crall JP, Stewart C V. Individual identification of the endangered Wyoming Toad *Anaxyrus baxteri* and implications for monitoring species recovery. J Herpetol. 2016;50: 44–49. doi:10.1670/14-155

38. Pike N. Using false discovery rates for multiple comparisons in ecology and evolution. Methods Ecol Evol. 2011;2: 278–282. doi:10.1111/j.2041-210X.2010.00061.x

39. Nemesházi E, Ujhegyi N, Mikó Z, Kásler A, Lente V, Bókony V. Dataset for: Photo-based individual identification is more reliable than visible implant elastomer tags or toe tipping in young agile frogs. Figshare. 2025. doi:10.6084/m9.figshare.29320697

40. Patel NG, Das A. Shot the spots: a reliable field method for individual identification of *Amolops formosus* (Anura, Ranidae). Herpetozoa. 2020;33: 7–15. doi:10.3897/HERPETOZOA.33.E47279

41. Kenyon N, Phillott AD, Ross AA. Temporal variation in dorsal patterns of juvenile green-eyed tree frogs, *Litoria genimaculata* (Anura: Hylidae). Herpetol Conserv Biol. 2010;5: 126–131.

42. Bardier C, Székely D, Augusto-Alves G, Matínez-Latorraca N, Schmidt BR, Cruickshank SS. Performance of visual vs. software-assisted photo-identification in mark-recapture studies: a case study examining different life stages of the Pacific Horned Frog (*Ceratophrys stolzmanni*). Amphibia-Reptilia. 2020;42: 17–28. doi:10.1163/15685381-BJA10025

43. Davis HP, Vancompernolle M, Dickens J. Effectiveness and reliability of photographic identification methods for identifying individuals of a cryptically patterned toad. Herpetol Conserv Biol. 2020;15: 204–211.

44. Brannelly LA, Berger L, Skerratt LF. Comparison of three widely used marking techniques for adult anuran species *Litoria verreauxii alpina*. Herpetol Conserv Biol. 2014;9: 428–435.

45. Lunghi E, Bruni G, Manenti R, Ficetola GF. Use of visible implant elastomer on two amphibians orders (Anura and Caudata): data on efficiency and reliability. Atti X Congr Naz Soc Herpetol Ital. 2015;15: 163–167.

46. Grant EHC. Visual implant elastomer mark retention through metamorphosis in amphibian larvae. J Wildl Manage. 2008;72: 1247–1252. doi:10.2193/2007-183

47. Cabot ML, Troan B V., Heugten KA Van, Schnellbacher RW, Smith D, Ridgley F, et al. Migration and histologic effects of visible implant elastomer (VIE) and passive integrated transponder (PIT) tags in the marine toad (*Rhinella marina*). Animals. 2021;11: 3255. doi:10.3390/ani11113255

48. Li C, Busquets R, Campos LC. Assessment of microplastics in freshwater systems: a review. Sci Total Environ. 2020;707: 135578. doi:10.1016/j.scitotenv.2019.135578

49. Grafe TU, Stewart MM, Lampert KP, Rödel M. Putting toe clipping into perspective: a viable method for marking anurans. J Herpetol. 2011;45: 28–35.

50. Waddle JH, Rice KG, Mazzotti FJ, Percival HF. Modeling the effect of toe clipping on treefrog survival: Beyond the return rate. J Herpetol. 2008;42: 467–473. doi:10.1670/07-265.1

