## Supplementary material for "Photo-based individual identification is more reliable than visible implant elastomer tags or toe-tipping in young agile frogs": S1 Appendix

**S1 Appendix: Details of rearing conditions and exclusions from the experiment**

We collected agile frog eggs from 12 egg masses (47.551195°N, 18.926682°E) on 15 March 2023 for the purpose of a long-term experiment, which included a 6-day heat treatment during the larval stage (results of the treatment will be discussed elsewhere). Twelve tadpoles were haphazardly chosen from each sampled egg mass when they reached the swimming stage (developmental stage 25 as defined by Gosner [1]), and were housed individually under laboratory conditions until the end of the treatment. The remaining tadpoles were released at their site of collection. In the laboratory, temperature was set to 16°C for the eggs and it gradually increased to c.a. 18°C for tadpoles and artificial light followed the natural diurnal cycle. During the course of the 6-day heat treatment, tadpoles in the control group were kept at 18.2°C ± 0.5°C (mean ± SD) and those in the treatment group were kept at 28.15°C ± 0.45°C (mean ± SD). Until the end of the treatment, the tadpoles were housed individually in white plastic boxes containing reconstituted soft water (RSW; reverse-osmosis filtered, UV-sterilized, aerated and reconstituted tap water). RSW volume in the individual boxes was 1 L for two weeks from developmental stage 25 onwards, and we increased it to 1.7 L for the treatment period. Twice a week, we changed the rearing water and fed the tadpoles with slightly boiled chopped spinach.

Subsequently to the treatment, sibling groups of heat-treated and control tadpoles were kept in separate outdoor mesocosms (i.e. six tadpoles per container; mesocosms were set up as described in [2]) until the start of metamorphosis. When an animal reached developmental stage 42 (appearance of forelimbs [1]), we assigned an individual identifier (Animal ID) to it as well as captured photographs of it (either by digital camera or smart phone), and we kept the individuals in separate boxes until tagging. During the peak of metamorphosis (between developmental stages 42 and 45 [1]) the animals were kept individually in tilted boxes that contained both shallow water and dry land. Individuals started metamorphosis between 29 May and 21 June, and completed it between 2 and 25 June. Thereafter, we kept the frogs outdoors in a shaded area, where they were kept individually until tagging, and were housed in sibling groups of six (three heat-treated and three control individuals) in 45-L translucent plastic housing boxes (56 × 39 × 28 cm) after tagging. We initially filled the housing boxes with a 1:1 mixture of heat-sterilized soil and sand, covered by pre-dried leaf litter and fitted with a small water bath. The insides of the boxes were sprinkled with aged tap water twice a week. The frogs were fed with springtails and crickets, except for the winter during which they hibernated in a climate chamber. For carrying out hibernation we followed the methods of [3], except that we housed the animals in groups of 4-6 siblings at c.a. 3.5°C ± 0.36 SD, and simulated the end of autumn and start of spring by gradually changing the temperature over the course of seven days from 10°C to 3.5°C and back, respectively.

Mortality rate was exceptionally high in two families from the beginning of the experiment: out of 12 individuals per egg mass, only three and four individuals survived until individual tagging, respectively, therefore these families were excluded from this study. However, out of the four animals still alive at tagging in the latter excluded family, one individual was accidentally assigned to another (non-excluded) family at start of metamorphosis (we realised this mistake because 13 instead of 12 animals were assigned to the family). Because the identity of the mistakenly assigned individual remained unknown, that individual stayed in the experiment. The froglets from the families that we excluded from the statistical analyses of this study were marked and kept in a single box like the rest of the animals, but they all died before the first hibernation, and we included their dissection data in tallying the occurrence of VIE material in internal organs. Additionally, we excluded two frogs from the analyses because they were inadvertently marked with two VIE tags instead of one (these were not counted in the N=113 in our study). The housing groups of frogs were never rearranged (i.e. when individuals died in a housing box, the reduced-size group was kept as it was).

### References


1. Gosner KL. A simplified table for staging anuran embryos and larvae with notes on identification. Herpetologica. 1960;16: 183–190. doi:10.2307/3890061

2. Bókony V, Üveges B, Móricz ÁM, Hettyey A. Competition induces increased toxin production in toad larvae without allelopathic effects on heterospecific tadpoles. Funct Ecol. 2018;32: 667–675. doi:10.1111/1365-2435.12994

3. Kásler A, Holly D, Herczeg D, Ujszegi J, Hettyey A. Chytridiomycosis and climate change: exposure to *Batrachochytrium dendrobatidis* and mild winter conditions do not increase mortality in juvenile agile frogs during hibernation. Anim Conserv. 2023;26: 654–662. doi:10.1111/acv.12851
