## Supplementary material for "Photo-based individual identification is more reliable than visible implant elastomer tags or toe-tipping in young agile frogs": S2 Appendix

**S2 Appendix: Supplementary figures.**

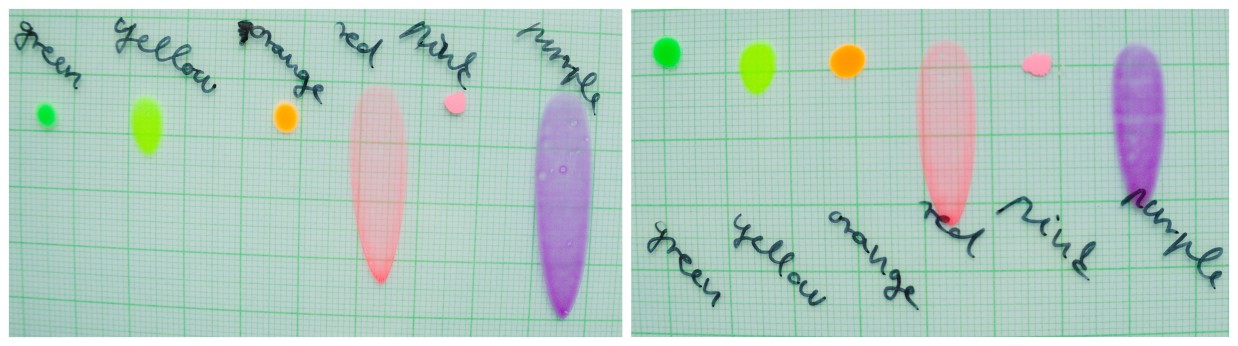

Fig S1. Self-made VIE reference cards prepared via lamination of the injected elastomers**.** Because the applied amount of VIE material of each colour on each card was similar, long smear systematically produced by the lamination process for red and purple indicates lower viscosity of these chemical mixtures. This was assumably caused by unintentional application of uneven volumes of VIE components (coloured elastomer base and curing agent) in the mixture that we prepared before injection.

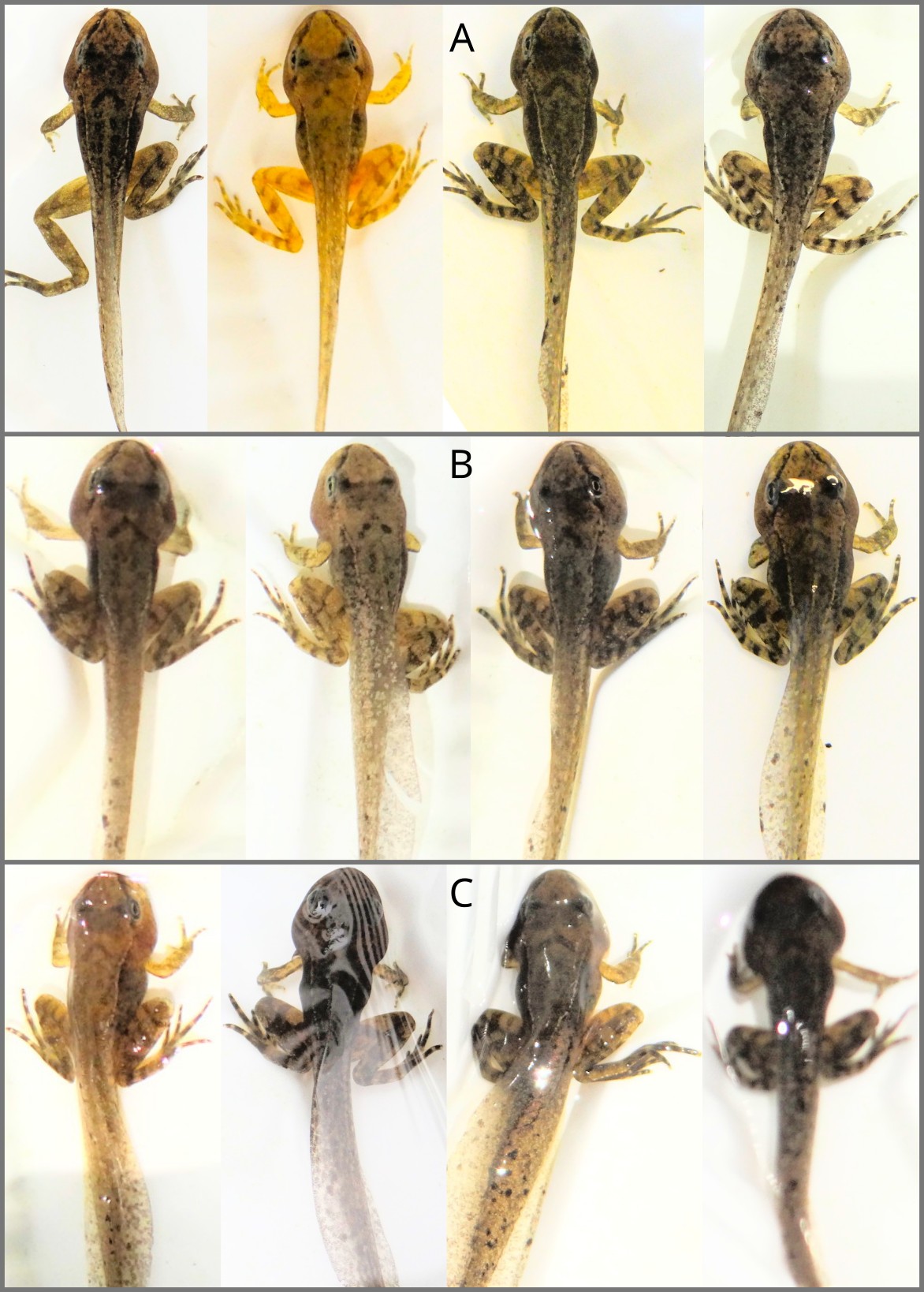

Fig S2. Quality categories used for photos captured at metamorphosis**.** According to the visibility of patterns across the back and legs, images were categorized as good (A), medium (B) or poor (C). Note that pattern visibility was influenced by posture, photo angle, focus and glare. Exposure curves were applied for each image to facilitate pattern visibility.

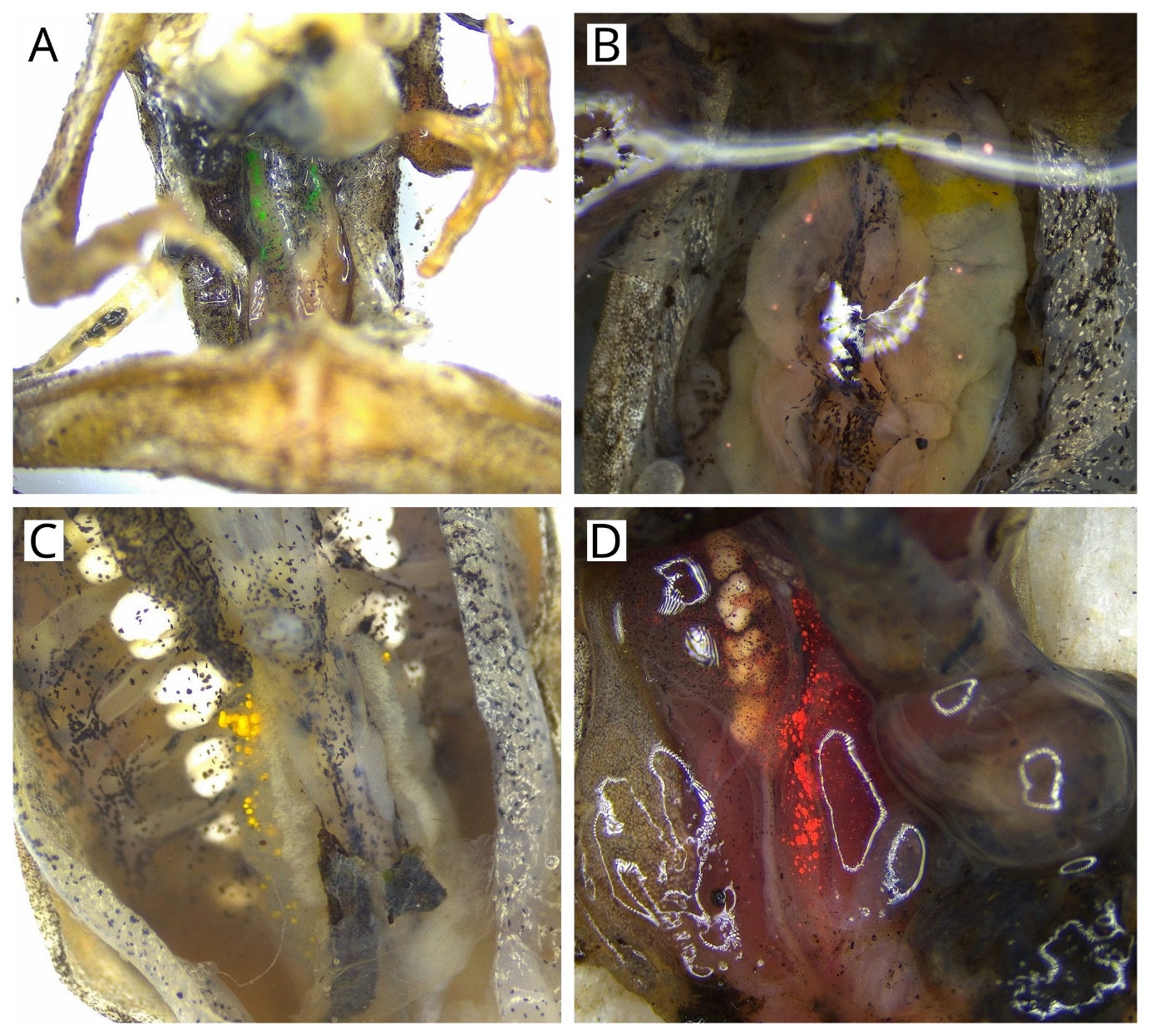

Fig S3. Examples of dissected juvenile agile frogs where VIE migrated into internal organs**.** Green (A), pink (B), orange (C), and red (D) VIE material is shown in the kidneys and the serous membranes (the liver and gastrointestinal tract were removed before taking the pictures) 62, 12, 380, and 415 days after tag injection. Taking into account froglets that died in this study as well as froglets that died in two families that we excluded from the study due to high mortality before marking (but they were marked for other purposes), we noticed VIE material in the internal organs for all six colours.

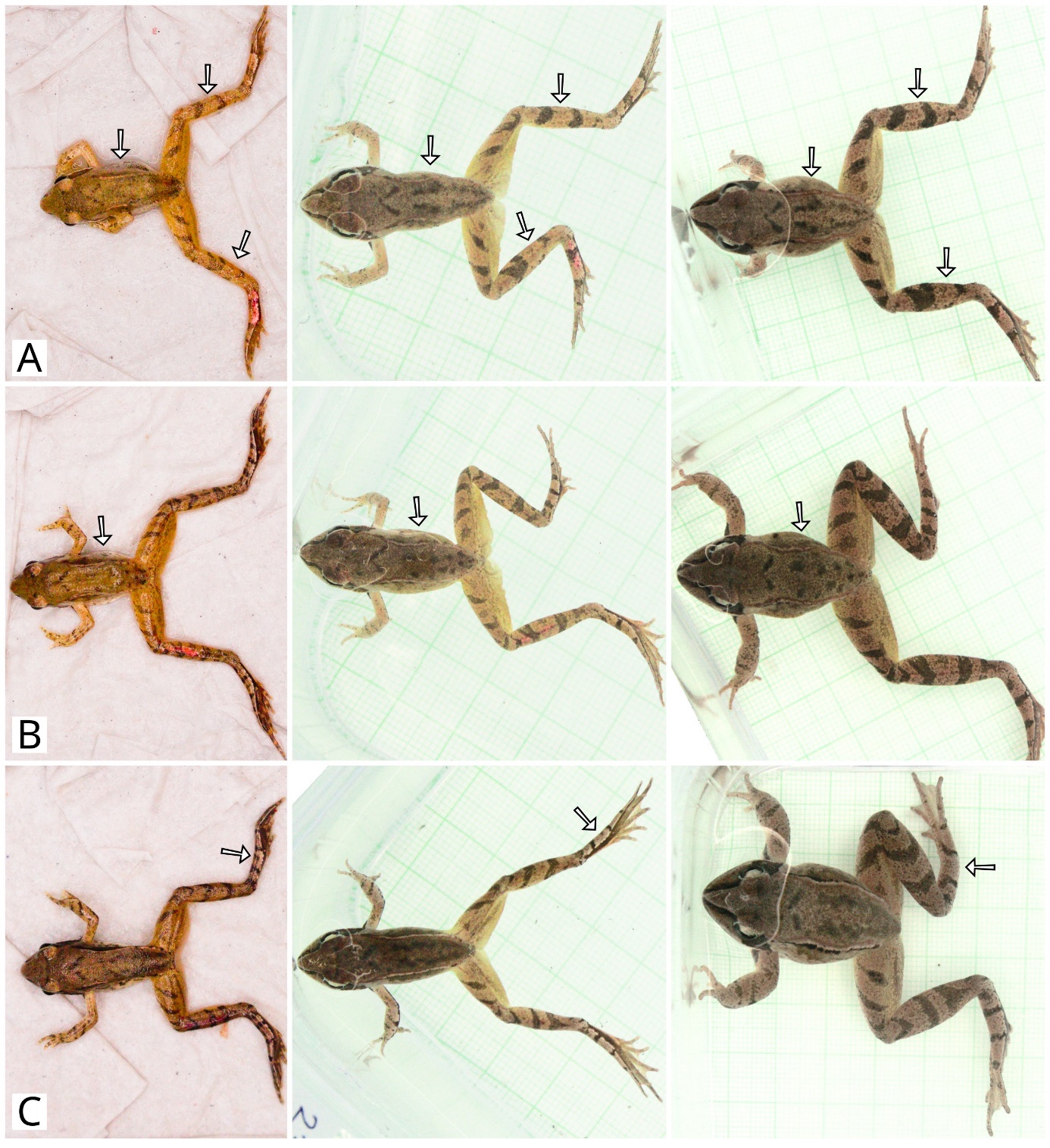

Fig S4. Examples of individuals where changes were detected in melanin-based patterns on the back or legs**.** Images within each row (A-C) feature the same individual at tagging in 2023 July, at the first checking event in August 2023 and at termination of the experiment in 2024 November, from the left to the right, respectively. Arrows indicate the body parts where changes were detected: melanisation increased over time (A, B) or a pattern disappeared (C). Note that apparent differences on the left shin on panel C were caused by gradual disappearance of the purple VIE tag. Exposure curves were applied for each image to facilitate pattern visibility.

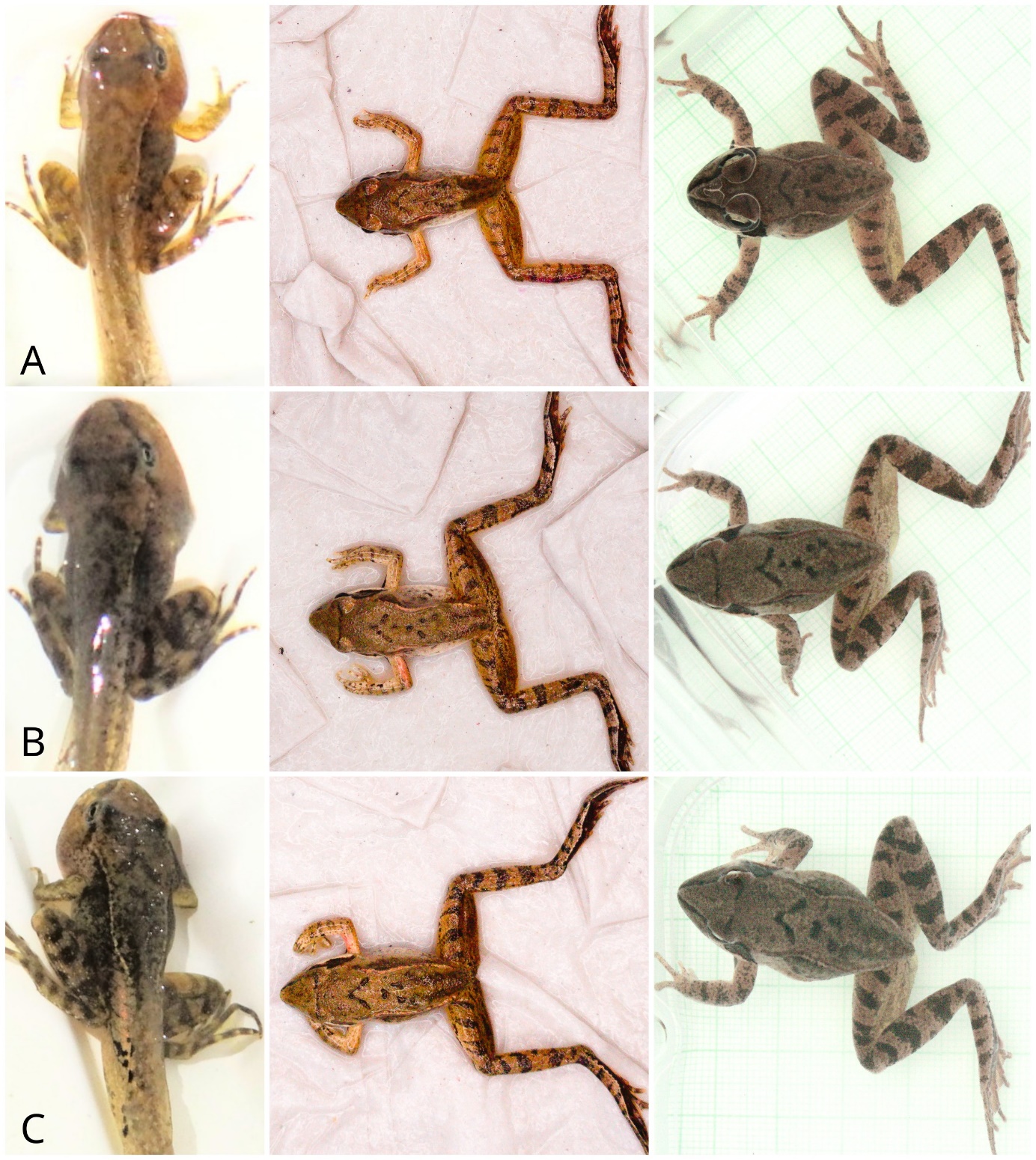

Fig S5. Examples of stable melanin-based individual patterns across the experiment**.** Images within each row feature the same individual (A-C) at metamorphosis in 2023 May-June, at tagging in 2023 July and at termination of the experiment in 2024 November, from the left to the right, respectively. Note that most leg stripes were already present at metamorphosis, and overall melanin-pattern distribution across the body remained stable after reaching the froglet stage. Individuals on panels B and C of this figure are siblings and B was mistakenly identified as C on-site in November 2024. Photographs clarified the true identity of the individual. Exposure curves were applied for each image to facilitate pattern visibility.
